# Novel Mechanism for Surface Layer Shedding and Regenerating in Bacteria Exposed to Metal-Contaminated Conditions

**DOI:** 10.1101/457358

**Authors:** Archjana Chandramohan, Elodie Duprat, Laurent Remusat, Severine Zirah, Carine Lombard, Kish Adrienne

## Abstract

Surface layers (S-layers) are self-assembling, ordered structures composed of repeating protein subunits found as components of the cell walls throughout the Bacteria and the Archaea. S-layers act as an interface between prokaryotic cells and their surrounding environment, and provide protection for microorganisms against diverse environmental stresses including heavy metal stress. We have previously characterized the process by which S-layers serve as a nucleation site for metal mineralization in the presence of high concentration of metals. Here, we test the hypothesis originally proposed in cyanobacteria that a “shedding” mechanism exists in prokaryotes for replacing S-layers that have become mineral-encrusted. We used a metallotolerant gram-positive bacterium bearing an S-layer, *Lysinibacillus* sp. TchIII 20n38, as a model organism. We characterize for the first time a mechanism for resistance to metals through S-layer shedding and regeneration. S-layers nucleate the formation of Fe-mineral on the cell surface, leading to the encrustation of the S-layer. Using a combination of scanning electron microscopy (SEM) and nanoSIMS, we show that mineral-encrusted S-layers are shed by the bacterial cells, and the emerging cells regenerate new S-layers as part of their cell wall structure. This novel mechanism for the survival of prokaryotes in metal-contaminated environments may also provide elements necessary for the development of renewable systems for metal bioremediation.

## 1 Introduction

Environmental contamination by metals and radionuclides from activities such as mining and nuclear power generation pose a serious risk to human health. The sudden, accidental release of high concentrations of iron from acid mine drainage from the Gold King Mine polluted the Animas River in 2015, mixing downstream with phosphates from agricultural runoff (Rodriguez-Freire et al.,2016). Metal contamination affected both water supplies from soluble metals, and sediments after the sedimentation of the majority of released metals. Sampling of soils contaminated by metals and radionuclides near the former Chernobyl nuclear reactor site (Chapon et al., 2012) and in uranium mining waste piles in Germany (Pollmann et al., 2006) have identified bacteria of the genre *Lysinibacillus* tolerant to these contaminants. *Lysinibacillus* formerly classified as part of the *Bacillus* genre (Ahmed et al., 2007)) gram-positive bacteria, with a peptidoglycan cell wall enclosed by a surface layer (“S-layer”) attached non-covalently to the lipopolysaccharides of the outer membrane (reviewed in (Sleytr et al., 2014)). These S-layers have proven to be a key mechanism for metallotolerance in *Lysinibacillus* as they have been shown to bind U, Pd(II), Cu, Pt(II), and Au(III) (Pollmann et al., 2006).

S-layers, however are not unique to *Lysinibacillus.* They are common components of the cell envelopes of both bacteria and archaea. S-layers are formed by self-assembly of repeated protein monomers into ordered structures (oblique, square, or hexagonal) depending on the number of subunits composing the ordered structure. This self-assembly occurs even in the absence of cells *in vitro*; a capacity has been exploited in biotechnology in everything from the development of vaccine to nanomaterials to filtration technologies (Sleytr et al., 2011).

S-layers form the interaction interface between prokaryotic cells and their external environment, and are therefore in contact with metals and other ions present. Nucleation of mineralization by S-layers was first noted in cyanobacteria by Schultze-Lam in the early 1990s (Schultze-Lam et al., 1992). Cyanobacterial S-layers were demonstrated to nucleate the formation of carbonates of calcium, magnesium, and strontium (Schultze-Lam and Beveridge, 1994). In their 1992 article, Schutlze-Lam proposed the hypothesis that mineral-encrusted S-layers are shed from cyanobacteria as part of a protective mechanism to ensure that essential cell activities are maintained despite cell wall mineralization (Schultze-Lam et al., 1992). This hypothesis was, however, never fully tested. Since then, S-layer nucleation of mineralization has been observed in a range of bacteria (Konhauser et al., 1994; Phoenix et al., 2000) and archaea (Kish et al., 2016).

S-layer mineralization is a mechanism for metallotolerance with potential application in bioremediation. S-layer shedding and regeneration in metallotolerant bacteria already inhabiting contaminated environments provides an added benefit of being a renewable system. Here, we describe the shedding and regeneration of mineral-encrusted S-layers in the metallotolerant environmental isolate *Lysinibacillus* sp. TchIII 20n38.

## 2 Materials and Methods

### 2.1 Culture and growth conditions

The bacterial strain used was an environmental strain isolated in 2009 from soils near a radionuclide-contaminated site (Chapon et al., 2012). This strain, referenced as *Lysinibacillus* sp. TchIII 20n38, was cultured at 30 °C in Luria Bertani (LB) medium under aerobic conditions with agitation (180 rpm) to mid-exponential, late-exponential, and stationary growth phases (OD_600nm_ = 0.3, 0.6, and 1.0, respectively). The culture medium was then removed and the cells washed in MilliQ-H_2_O by gentle centrifugation (2600 x g, 15 min, room temperature). In order to determine the mechanisms of resistance of *Lysinibacillus* sp. TchIII 20n38 cells to the presence of heavy metals, the cells were resuspended to an equivalent cell density in a Fe-rich solution at a similar pH to that found in the Chernobyl isolation (10 mM NaH_2_PO_4_, 10 mM FeSO_4_, pH=4.5), and agitated (150 rpm, 30 °C) with for up to 5 days. Cells were filtered and observed by scanning electron microscopy as described below.

### 2.2 Mineralization Recovery Time Course

In order to test the hypothesis that mineral-encrusted S-layers are shed and regenerated, a time course of recovery was followed after Fe-mineralization as follows: *Lysinibacillus* sp. TchIII 20n38 cells were grown to mid-exponential growth phase (OD_600nm_ = 0.3) in LB (30 °C, 180 rpm). The culture medium was then removed and the cells washed in MilliQ-H_2_O by gentle centrifugation (2600 x g,15 min, room temperature). The cells were then resuspended to an equivalent cell density in either a mineralization solution (10 mM NaH_2_PO_4_, 10 mM FeSO_4_, pH=4.5), or a nutrient-free buffered solution at the same pH (10 mM NaH_2_PO_4_, pH=4.5), and agitated (30 °C, 150 rpm) for 16 h. The mineralization solution was then removed and the cells washed in MilliQ-H_2_O by gentle centrifugation (2600 x g, 15 min, room temperature), and replaced by growth medium (LB, or labelled growth medium).

For NanoSIMS experiments, LB was replaced with a defined growth medium approximating LB but containing either ^14^N or ^15^N (Celtone^®^, Cambridge Isotopes, USA). While specified for use with bacteria for isotope analyses, this media was found to contain inhibitory concentrations of trace metals, which we were able to remove by precipitation by addition of an excess of buffered phosphate (10 mM) over 24 h with agitation at 150rpm, followed by filtration using a 0.22 p,m filter. This metal-depleted medium containing 5 g/L Celtone^®^ was completed with 5 g/L acetate as a C-source, and basal salts (15 mM ammonium sulfate, 0.2 mM MgSO_4_,17.6 mM KH_2_PO_4_, 32.7 mM NaH_2_PO_4_), as determined by our preliminary optimized experiments with *Lysinibacillus* sp. TchIII 20n38. In addition, cultures grown in the presence of ^14^N rather than^15^N rapidly ceased vegetative growth and sporulated. Therefore instead of a standard labelling medium composed of both ^14^N and ^15^N, a 100% ^15^N-labeled medium, was used to follow the time course of recovery after Fe-exposure. An additional culture was resuspended in a 100% ^14^N medium and immediately sampled as a baseline control for N isotope abundances. Cultures were incubated in the ^15^N-labeled medium at 30 °C with agitation (180 rpm) over a time course of recovery, for both mineralized (M) and nonmineralized (NM) cultures. Aliquots were removed immediately after addition of growth medium (T0), and then every 24 h (1d, 2d). At each time point, approximately 20mL aliquots were removed for optical density measurements, optical microscopy verification of cell morphology, and filtered for scanning electron microscopy observations and NanoSIMS analyses as described below. Abiotic (non-inoculated) controls were used for comparison to distinguish mineralization due to the presence of *Lysinibacillus* sp. TchIII 20n38 cells.

### 2.3 Scanning Electron Microscopy

Aliquots of bacterial cultures as well as abiotic controls (not inoculated with *Lysinibacillus* sp. TchIII 20n38 cells) were filtered through a 0.2 p,m GTTP isopore polycarbonate filters using a Swinnex filter holder (Merck Millipore, Darmstadt, Germany). Filters were then air-dried, mounted on aluminum supports with carbon tape, and coated with carbon (7-8 nm thickness), gold (7 nm thickness), or platinum (5 nm thickness). Scanning electron microscopy (SEM) observations were performed using two different instruments; a Hitachi SU 3500 SEM installed at the electron microscopy platform of the Muséum National d’Histoire Naturelle (Paris, France), and a Zeiss Ultra 55 field emission gun SEM equipped with a Brucker EDS QUANTAX detector (Brucker Corporation, Houston, TX, USA) installed at IMPMC (Sorbonne Université, Paris, France). For observations using the Hitachi SU 3500 instrument observations were made in secondary electron mode with an acceleration voltage of 15 kV. SEM-FEG images were acquired in secondary electron mode using with the Zeiss Ultra 55 instrument with an in column detector (InLens) at 2 kV to 5 kV and a working distance of 3 mm. Energy dispersive X-ray spectroscopy (EDX) analyses were performed at 15 kV and a working distance of 7.5 mm after calibration with reference copper.

### 2.4 Nano Secondary Ion Mass Spectrometry (NanoSIMS)

NanoSIMS sample preparations followed the protocol of (Miot et al., 2015). Briefly, aliquots of bacterial cultures sampled from labeled and unlabeled media filtered through 0.2 µm GTTP isopore polycarbonate filters previously Au-coated (20 nm thickness) using a Swinnex filter holder (Merck Millipore, Darmstadt, Germany). Quantitative ion images were recorded by the NanoSIMS50 (Cameca, Gennevilliers, France) installed at the National Museum of Natural History of Paris,France. All measurements were performed using the same analytical conditions. A Cs + primary beam of 0.8 pA scanned an area of 20 µm x 20 µm, divided into 256 pixels x 256 pixels, with a counting time of 1 ms per pixel. Secondary ion images of ^31^P^16^O-, ^12^C^14^N-, and ^12^C^15^N- were recorded. The mass resolution power was adjusted to 9000 to resolve isobaric interferences at mass 27 such as ^13^C^14^N- or ^11^B^16^O from ^12^C^15^N-. Before any analysis, the surface of each sample was pre-sputtered during 5 min with a 80 pA Cs- primary ion beam over 30 pm × 30 pm to eliminate the contamination of the surface, and reached the stable state of sputtering (Thomen et al., 2014). Instrument stability was verified throughout the session using a type 3 kerogen standard. NanoSIMS data were then processed using the IMAGE software (L. Nittler, Carnegie Institution for Science, Washington, DC, USA).

### 2.5 Statistical Analyses

The preference of this strain of *Lysinibacillus* for ^15^N to maintain vegetative growth eliminated the possibility of using a ^14^N control throughout the time course of recovery. In order to automatically remove the random noise from all the ^12^C^14^N- and ^12^C^15^N- NanoSIMS images, we defined ^12^C^14^N- and ^12^C^15^N- independent thresholds based on their respective distribution for the images of samples that were resuspended in the ^14^N-labelled medium. Each elemental distribution was fitted by two Gaussian components (R-package mixtools, (Benaglia et al., 2009)). We define threshold as the mean of the 97.5th percentile of the first Gaussian component (noise) and the 2.5th percentile of the second one (signal).

For each image, the denoised dataset further used for statistical analyses is composed only of pixels with both ^12^C^14^N- and ^12^C^15^N- values above the respective thresholds. The isotope abundance ^12^C^15^N- / (^12^C^14^N- + ^12^C^15^N-) of this dataset (hereafter named processed ^15^N/(^14^N+^15^N) ratio) was used to follow the kinetics of incorporation of N (as part of protein production) by non-mineralized and mineralized bacteria. For each image, the distribution of processed ^15^N / (^14^N + ^15^N) ratio was fitted using Gaussian mixture modeling (R-package mclust, (Scrucca et al., 2016)) in order to infer subpopulations of pixels. The best univariate model, composed of *k* Gaussian components with either equal or unequal variance, was selected based on Bayesian Information Criterion. The mixing proportions for the components represent the proportions of the *k* subpopulations of pixels.

Cluster analysis was performed in order to group the samples according to their subpopulation composition. Each image was described by a vector of 10 values, each corresponding to the sum of the mixing proportions for the Gaussian components whose mean falls in a given ^15^N/(^14^N+^15^N) ratio interval (10 intervals of size 0.1 each). An image-to-image distance matrix generated by computing Bray-Curtis dissimilarity index between all the pairs of vectors was used for hierarchical agglomerative clustering of images (unweighted pair group method with arithmetic mean linkage).

All analyses were conducted in R version 3.2.3 (R Core team, 2015).

## 3. Results

### 3.1 *Lysinibacillus* sp. S-layers Become Encrusted After Exposure to Iron

*Lysinibacillus* sp. TchIII 20n38 was isolated from soils contaminated by radionuclides and metals, resulting in a moderately acidic pH (5.5) (Chapon et al., 2012). Our work with this strain, like other isolates from the same site, has shown that it is resistant to a range of heavy metals and radionuclides (*article in preparation*). To determine the mechanisms of this metallotolerance, we exposed cells to a Fe-rich solution at acidic pH in the absence of the preferred carbon source for this strain, acetate, while maintaining high levels of phosphate required by *Lysinibacillus* sp.. Non-metabolic metal-tolerance mechanisms are favored under these conditions.

After exposure, Fe-minerals were observed to form on the surface of *Lysinibacillus* sp. cells leading to complete encrustation of the cells over time (see Fig. 1). EDX analyses of mineral-encrusted cells confirmed the composition as a Fe-phosphate (see Supplementary Figure 1). Some abiotically formed Fe-phosphates were also observed, which were easily distinguishable from mineralized S-layers as aggregates of larger spherically-shaped minerals not associated with cells, and matching the types of minerals observed in non-inoculated controls.

**Figure 1.**
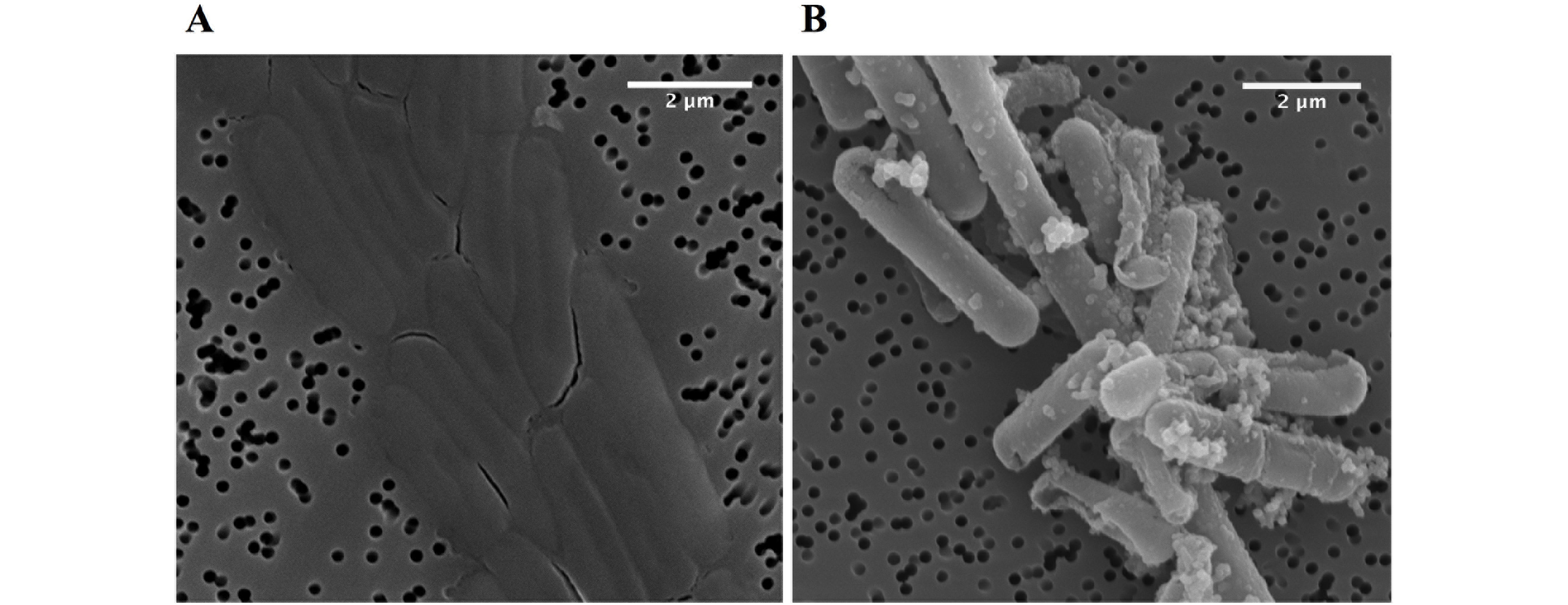
Scanning electron microscopy images in secondary electron mode of *Lysinibacillus* sp. TchIII 20n38 cells. **(A)** grown under optimal conditions, and **(B)** after exposure to an iron-rich solution (10 mM NaH_2_PO_4_, 10 mM FeSO_4_, pH=4.5). Scanning electron microscopy images acquired in secondary electron mode.

Cells were fully mineral encrusted after 16 h of exposure to the Fe-rich solution, whether cells were exposed in mid-exponential, late-exponential, or stationary growth phase (OD_600nm_ = 0.3, 0.6, and 1.0, respectively). Longer exposures (20 h and 41 h) did not alter the extent mineralization.

Attempts to confirm whether the mineralization observed on *Lysinibacillus* sp. TchIII 20n38 cells was due to a completely non-metabolic process, or whether active metabolism by living cells was necessary for S-layer mineralization were limited due to an inability to obtain dead cells without damaging the S-layer containing cell envelope. This despite multiple trials employing various antibiotics targeting non-cell envelope structures, and testing them over a large range of concentrations and durations [tetracycline (10-2000 p,g/mL for 1 h-5 d in LB, buffer, or MilliQ-H_2_O) ofloxacin (10-500 p,g/mL), and heat treatments (up to 55°C)]. The fact that *Lysinibacillus* sp. TchIII 20n38 cells grow optimally as heterotrophs without added metals suggests that the role of any metabolic processes in S-layer mineralization was secondary to non-metabolic processes.

### 3.2 Physiological State of Cells Depends on Times of Mineralization and Recovery

Replacement of mineral-encrusted cells into a rich growth medium demonstrated that *Lysinibacillus* sp. TchIII 20n38 cells were able to resume proper cell division after complete Fe-mineral encrustation. The exposure of mid-exponential growth phase cells (OD_600nm_= 0.3) to the Fe-rich solution for 16 h followed by recovery of the cultures in LB showed that cells resumed normal cell division (see Supplementary Figure 2). However, the physiological state of the cells during metal exposure affected the ability of cells to recover after mineral encrustation, resulting in various inhibitions of normal vegetative cell growth and division. For mid-exponential growth phase cells, exposures longer than 16 h to the Fe-rich solution resulted in the death of mineralized cells, and the formation of filaments by the small minority of cells observed without mineral-encrustation. In the case of both late-exponential growth phase and stationary phase cultures, even 16 hour-long exposures to Fe resulted in sporulation and/or cell death.

### 3.3 Mineral-Encrusted S-layers Can Be Shed

SEM observations were made of mid-exponential growth phase *Lysinibacillus* sp. TchIII 20n38 cells after a 16 h exposure to the Fe-rich solution. Mineralized S-layers devoid of a cell were observed, often with the cells located beside these empty mineralized S-layer shell (see Fig. 2 panel A, left and right images). After incubation in LB for up to 5 d following Fe-mineralization, cells were seen exiting mineralized S-layer cells, with cell division septa visible (see Fig. 2 panel B, top and bottom images), in concordance with increases in the optical density of the cultures (see Supplementary Figure 2).

### 3.4 *Lysinibacillus* sp. Cell Morphology Changes During S-layer Shedding

The cell morphology in both mineralized and non-mineralized cultures was altered over the time course of recovery (see Fig. 3). In non-mineralized cultures which were kept at the same pH as the mineralized cultures but in the absence of iron, cells gradually shrank in size and became ovoid in shape over two days of incubation in the metal-depleted Celtone^®^ medium. While cell death was minimal for non-mineralized cells, dead cells were easily distinguishable due to both their high ^12^C^14^N^−^ counts due to lack of ^15^N incorporation, and their sustained rod shape (see image of nonmineralized cells 1 d in Fig. 4). Non-mineralized cells also formed intracellular polyphosphate granules between 1 d and 2 d of incubation, as evidenced by analyses of ^31^P^16^O- counts (see Fig. 4).

**Figure 2.**
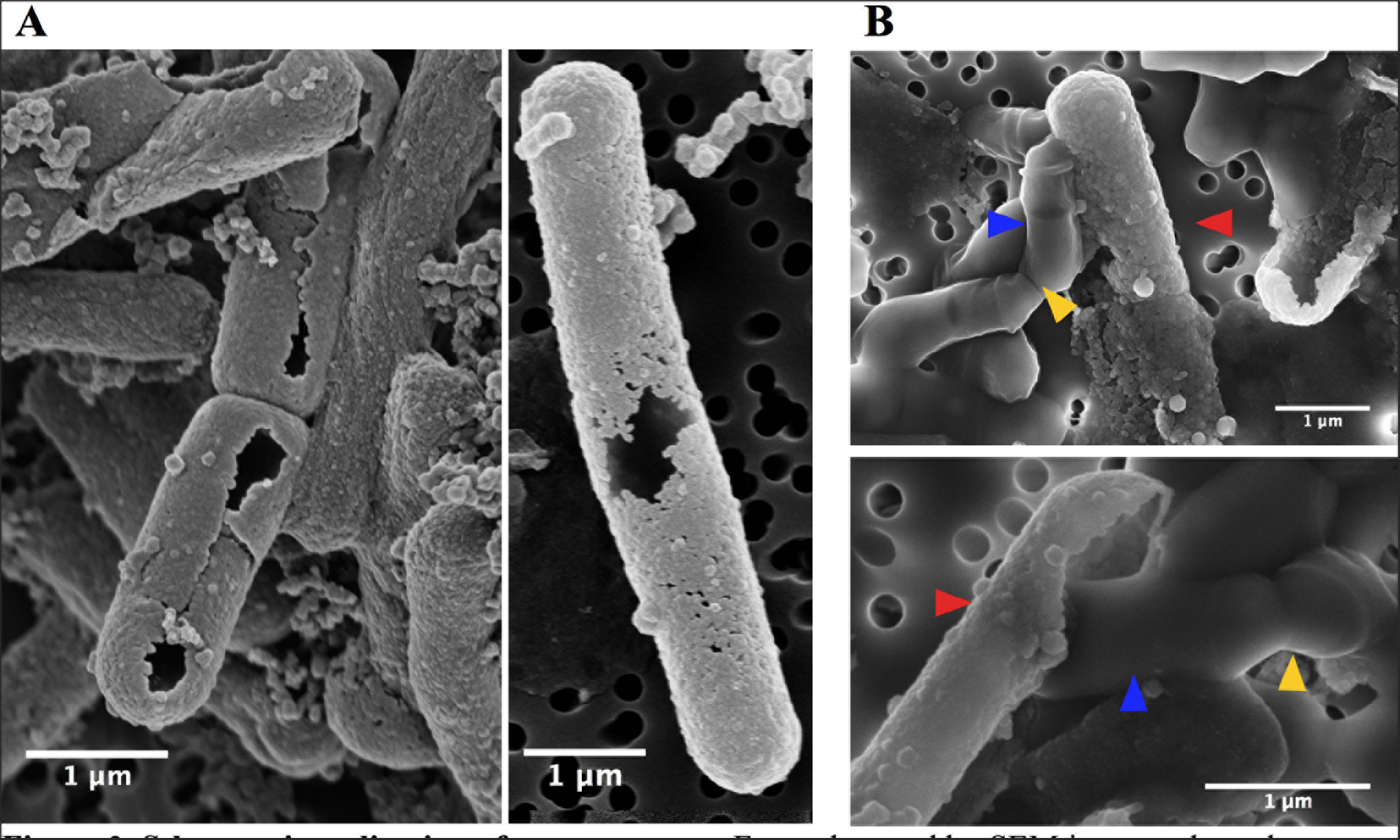
S-layer mineralization after exposure to Fe as observed by SEM in secondary electron mode. **(A)** Empty mineralized S-layer “shell” lacking a cell observed by SEM (secondary electron mode) after a 16 h exposure of a mid-exponential growth phase culture of Lysinibacillus sp. TchIII 20n38 to the Fe-rich solution. Empty S-layer mineral encrustation can be thick (lift-side image) or thinner and “lacy” (right-side image). **(B)** Top and bottom images show cells emerging from shed mineralized S-layers. Cells were in mid-exponential growth phase prior to exposed to Fe, followed by incubated in LB for up to 5 days (37 oC, 180 rpm). Blue arrows show emerging cells. Red arrows indicate mineralized S-layers. Yellow arrows indicate cell division septa.

**Figure 3.**
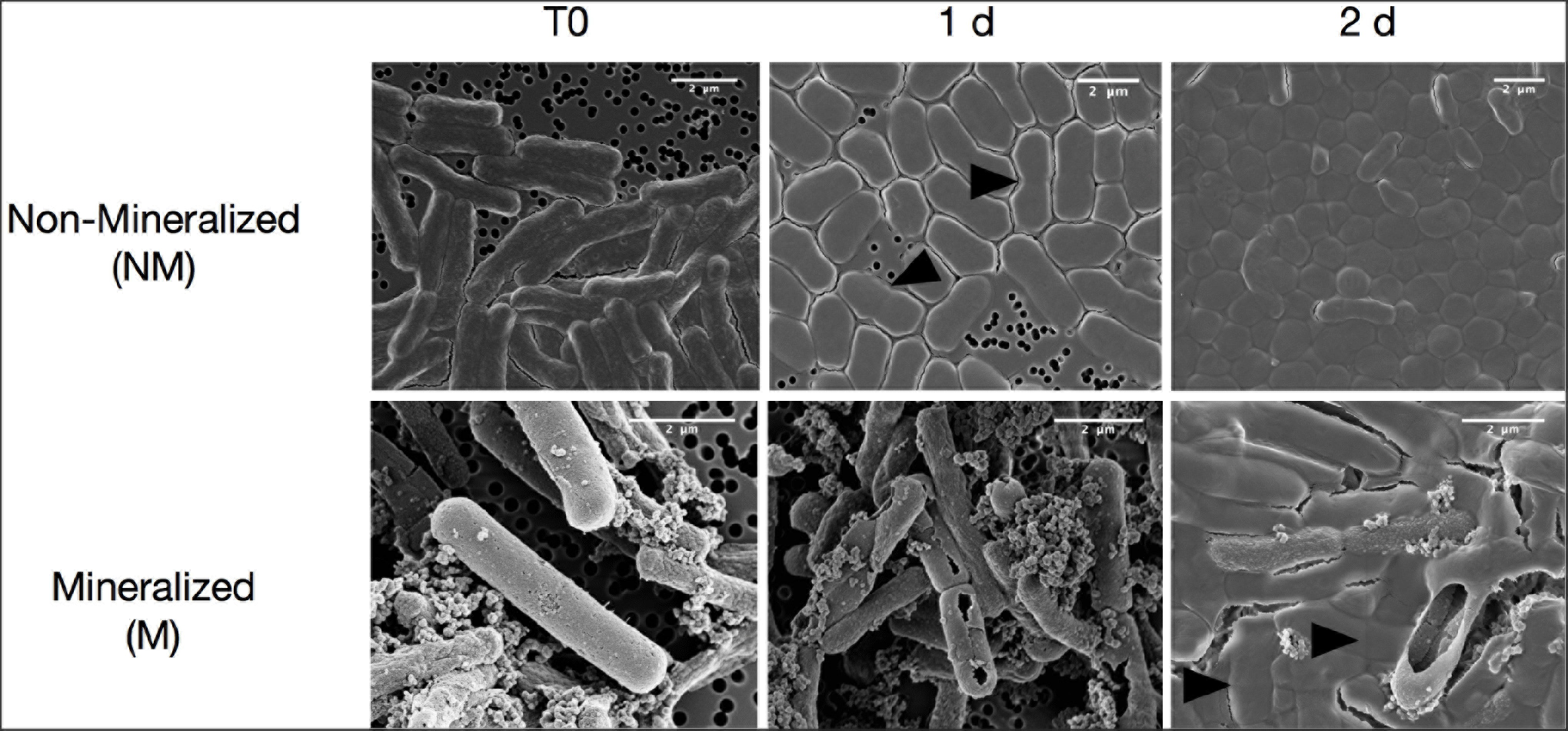
Scanning electron microscopy images of Lysinibacillus sp. TchIII 20n38 cells. (secondary electron mode) for mineralized and non-mineralized cells over a time course of recovery after Fe-exposure.

**Figure 4.**
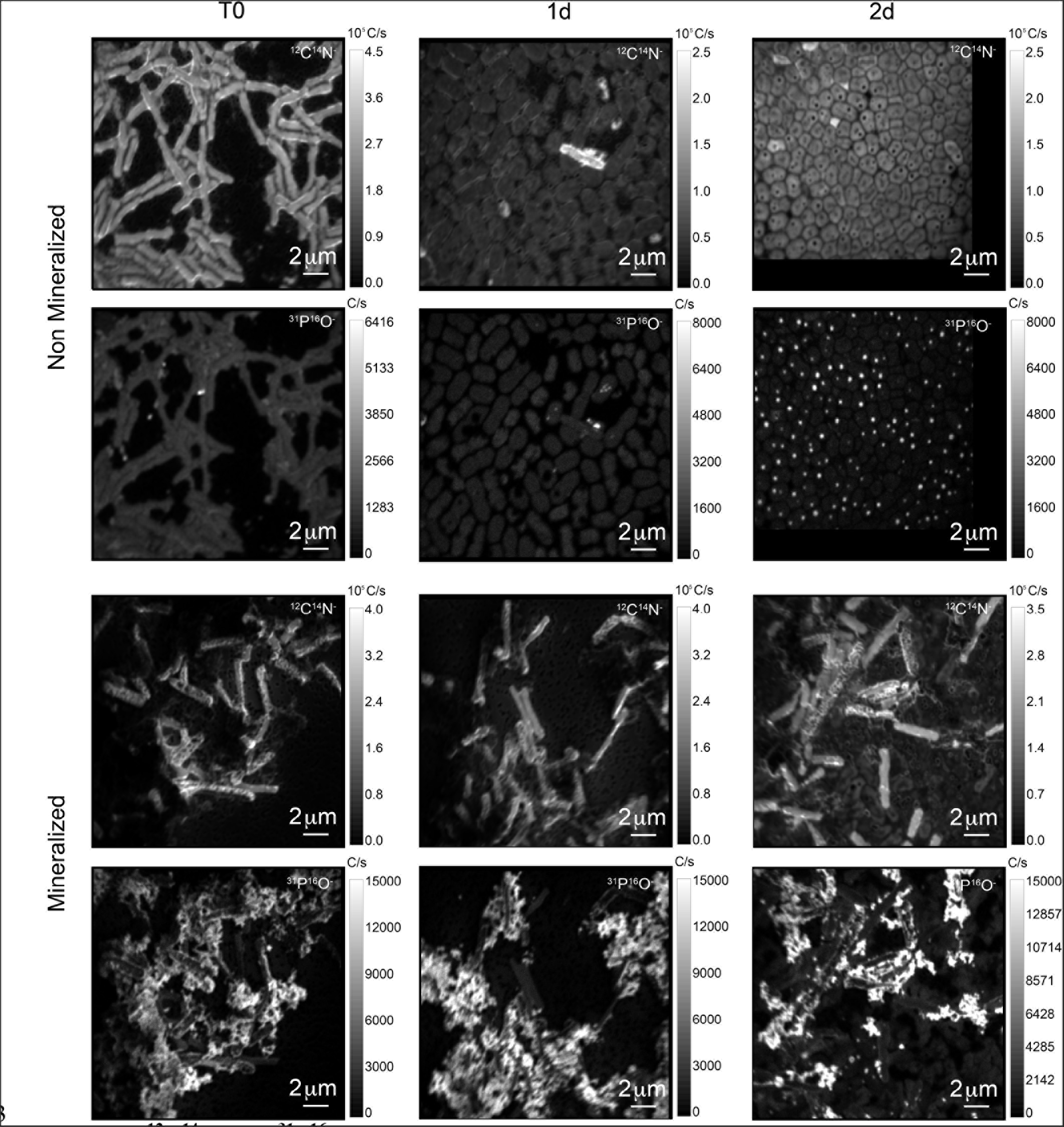
^12^C^14^N- and ^31^P^16^O- NanoSIMS images for both non-mineralized and mineralized cells over a time course of recovery after Fe-exposure.

Mineralized cultures showed little change over the first day of incubation. On the second day, however, cells were observed outside of their mineralized S-layer shells, with biofilm formation evidenced as a mucoid phenotype (see Fig. 3) together with increases in optical density of the cultures (see Supplementary Figure 3). SDS-PAGE and mass spectrometry confirmed the presence of S-layer glycoproteins throughout the time course of recovery (see Supplementary Figures 4 and 5). Shed, mineralized S-layers maintained the elongated rod shape of non-stressed *Lysinibacillus* sp. TchIII 20n38 cells and restricted ^31^P^16^O- to their surfaces (see Fig. 4), likely within the Fe-phosphate minerals analyzed by EDX in Figure 2. In comparison, newly emerged cells lacked surface phosphates, some cells concentrating phosphate as intracellular granules (see Fig. 4). On the third day of incubation, cells in both mineralized and non-mineralized cultures began to sporulate.

### 3.5 Sub-populations of Cells Co-exist During Time course of Recovery

In order to describe the process of S-layer regeneration after mineral encrustation, and to determine if the S-layers were indeed regenerated, we followed the recovery of *Lysinibacillus* sp. TchIII 20n38 cells over time after Fe-mineral encrustation. Both SEM observations of cell morphology and NanoSIMS analyses of cell activity using incorporation of nitrogen, needed for the production of new S-layer proteins. ^15^N-incorporation is an effective marker, as S-layer proteins are one of the most abundant cellular proteins, and roughly 20% of total protein synthesis can be dedicated to their production (Sleytr et al., 2007). The ^15^N incorporation over time was determined using ^12^C^15^N/(^12^C^15^N+^12^C^14^N) and statistical analyses of the ^15^N/(^14^N + ^15^N) were then performed (see example of sample M_T0_rep2 in Fig. 5, Supplementary Figure 6). In order to account for differences in cell morphology over time and between mineralized and non-mineralized samples, statistical analyses were performed using all pixels above the established threshold from each image. At least three processed ^15^N/(^14^N + ^15^N) images were analyzed per sample time point for samples incubated in medium containing ^15^N (two images each for natural abundance controls), with 10 to < 60 cells per image depending on the physiological state of the cells over the time course. Subpopulations of pixels were identified for each sample according to the distribution of processed ^15^N/(^14^N + ^15^N) ratio (see Fig. 5). Figure 5 shows that at the start of the time course (T0), the mineralized cells had ^15^N/(^14^N + ^15^N) ratios below 0.5 (panel A), with most cells having a ratio near 0.3 (panel B). Pixel subpopulations clustered around cells (panel C), with most mineralized cells and abiotic mineralization retaining a small amount of ^15^N (panel D) during the brief exposure prior to washing the cells in the first few minutes of the experiment.

**Figure 5.**
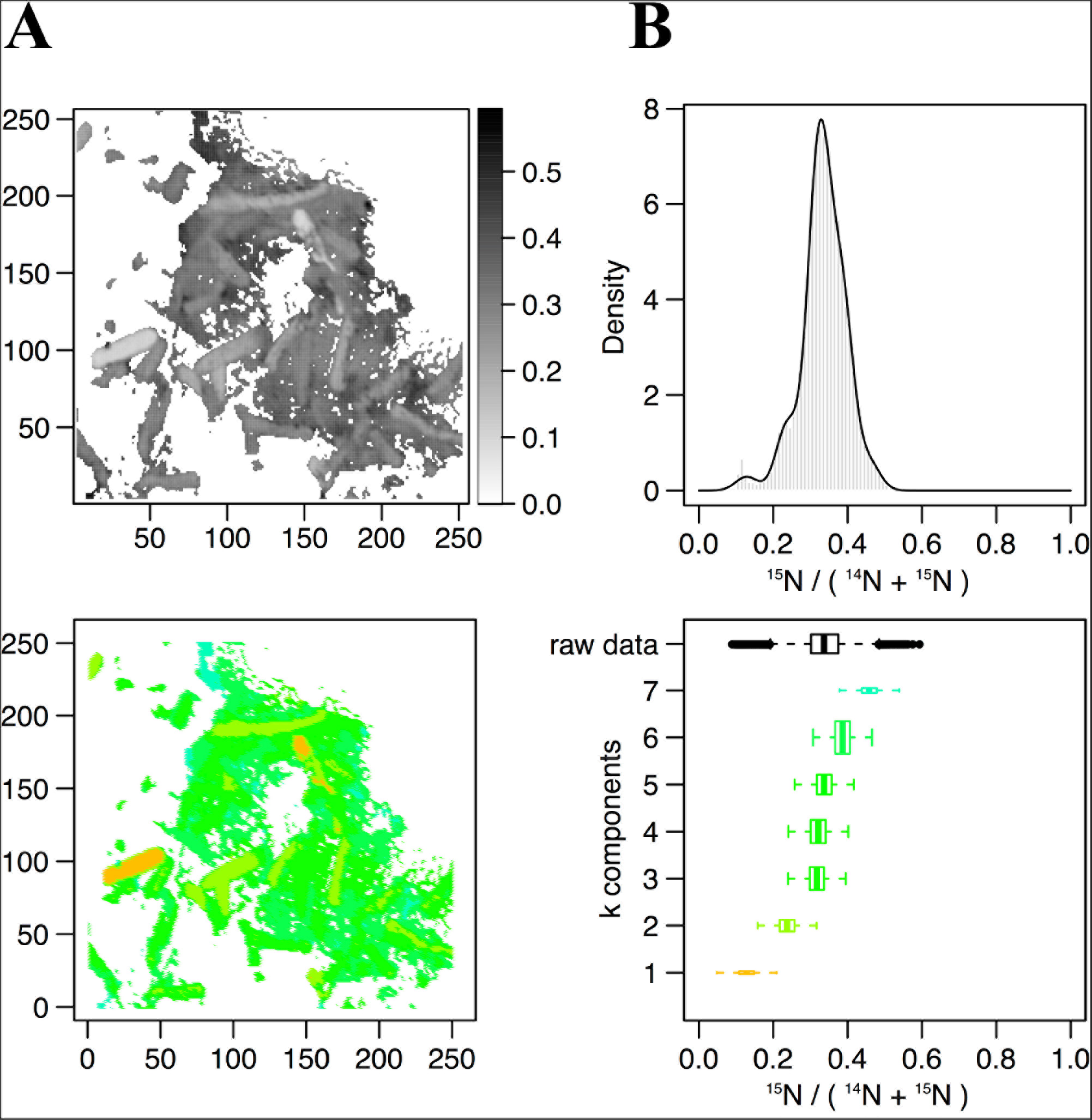
Statistical analysis of ^15^N/(^14^N^15^N) NanoSIMS images. Statistical analysis of ^15^N/(^14^N+^15^N) NanoSIMS image for mineralized cells after 2 days of recovery (M_2d_rep16). **(A)** Processed ^15^N/(^14^N+^15^N) ratio map. **(B)** Processed ^15^N/(^14^N+^15^N) ratio histogram (gray bars) and probability density (dark line) estimated by Gaussian mixture modeling. **(C)** Map of the pixel subpopulations (see (D) for color code). **(D)** Decomposition of the distribution of processed ^15^N/(^14^N+^15^N) ratios (labelled “raw data”) into 4 Gaussian components of unequal variance. The box boundaries indicate the first and third quartiles. The bold band inside the box corresponds to the median value. The horizontal dashed lines that extend from the box encompass the largest/smallest ratio values that fall within a distance of 1.5 times the box size from the nearest box hinge. If any, outliers are shown as individual points. Each component of the Gaussian mixture is represented by a random sample (n=10000) from the corresponding normal distribution. The box heights represent the proportions of pixel subpopulations (i.e. the mixing proportions for the components). A unique color scale is used for all images, the subpopulation color depending on the estimated mean of the Gaussian component.

Clustering of the NanoSIMS images according to their distribution of processed ^15^N/(^14^N + ^15^N) ratio showed that the samples tended to cluster over the experimental time course into three different groups; “natural abundance” (^14^N) T0 controls, low ^15^N incorporation (NM_T0, M_T0, M_1d), and significant ^15^N incorporation (NM_1d, NM_2d, M_2d) (see Fig. 6). The fact that controls for the natural abundance (NM_no incubation, M_no incubation) grouped separately from the T0 samples shows that even brief exposure to the ^15^N-labelled medium during cell resuspension had an effect on the isotopic composition of the cells. The T0 samples were therefore used as the baseline of comparison for all later time-points. The two remaining groups were composed of cells cultivated in ^15^N-labelled medium, grouped by whether or not cell division had restarted. Low ^15^N-incorporation samples (M_T0, NM_T0, M_1d) corresponded to cells not yet showing evidence of cell division, whereas samples with significant ^15^N incorporation (NM_1d, NM_2d, M_2d) showed clear evidence of active cell division when observed by SEM (see Fig. 3, black arrowheads) and measurements of optical density (see Supplementary Figure 3).

**Figure 6.**
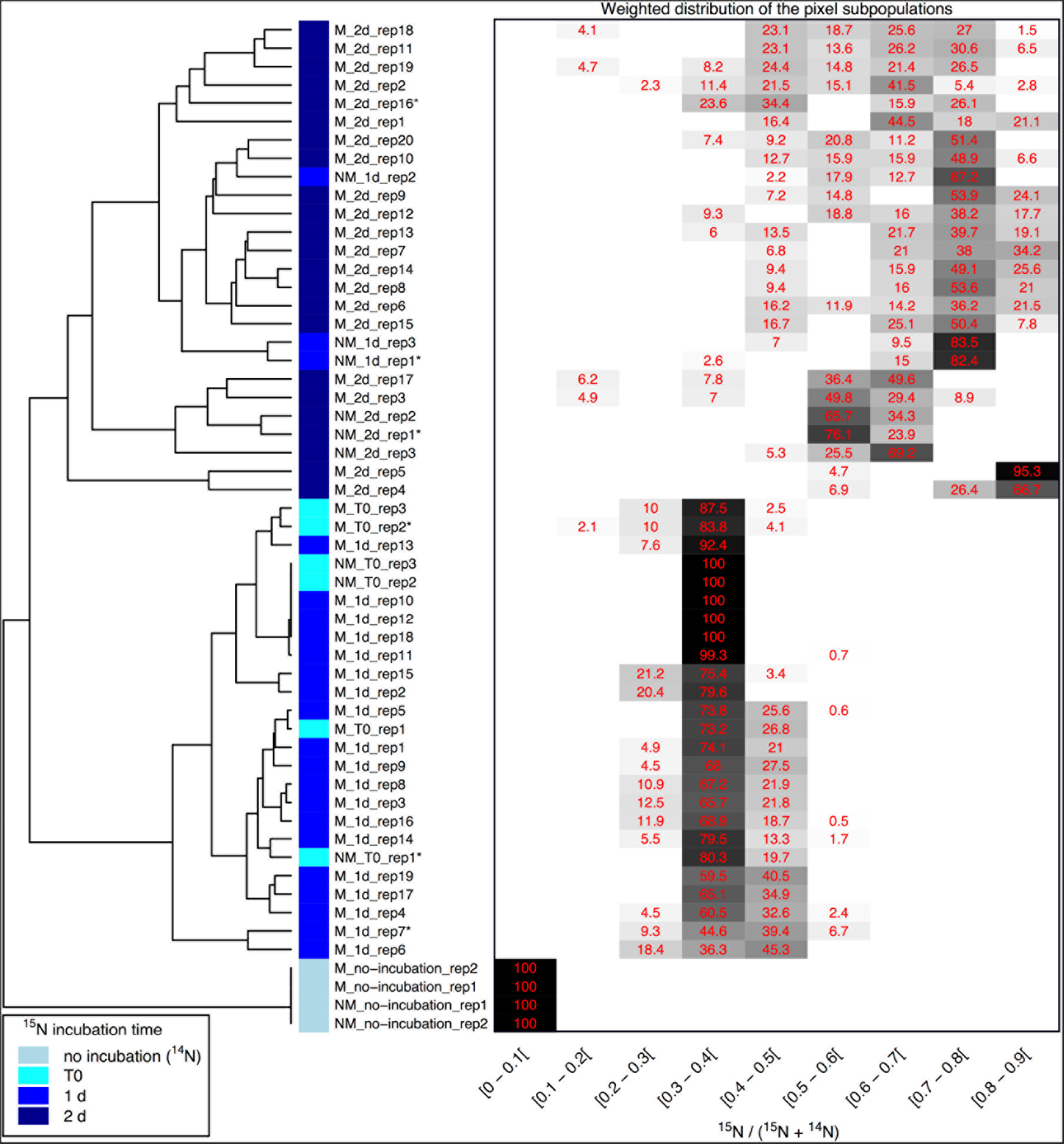
Clustering of ^15^N/(^14^N+^15^N) NanoSIMS images according to the distribution of pixel subpopulations. Each row of the heatmap (drawn on the right side of the figure) shows proportion (in %) of the pixel subpopulations for a single NanoSIMS image, and columns show the ^15^N/(^14^N+^15^N) ratio intervals. Each value corresponds to the sum of the mixing proportions for the Gaussian components whose mean falls in the given interval. Null proportions are hidden. Color scale ranges between white (0 %) and black (100 %). Each image is labelled according to the culture medium (either M or NM for Mineralized or Non-Mineralized, respectively), the ^15^N incubation time, and the replicate number (a NanoSIMS image numbering for a given sample). Each image is depicted by a color code according to the ^15^N incubation time of the sample (see color legend at bottom left). The heatmap rows are ordered according to the hierarchical clustering of images illustrated on the left side of the figure. Image labels ending with a star a re further illustrated in Figure 6 and Supplementary Figure 6.

Variations between replicate images were minimal for all samples with the exception of both mineralized and non-mineralized 2 d samples, as seen in the heat map representation. The weighted distribution of subpopulations remained low for all samples prior to restart of cell division (see Fig. 6 and Supplementary Figure 6, samples M_T0, M_1d, NM_T0). The increase in pixel subpopulations for samples showing signs of recovery and cell division (samples NM_1d, NM_2d, M_2d) reflects the heterogeneity of cell recovery. Heterogeneity in ^15^N incorporation was highest for cells immediately after S-layer shedding (M_2d samples), which is likely a reflection of natural variations in the capacity of *Lysinibacillus* sp. TchIII 20n38 cells to respond to stress.

### 3.6 S-layers are Regenerated Within Two Days of Fe-Mineral Encrustation

Some cell-free mineralized S-layer shells were observed by SEM immediately after 16 h exposure to the Fe-rich solution (see Fig. 3). However, the morphology of the cells remained generally unchanged through 1 d of incubation in the defined medium. At 2 d of incubation in this medium, SEM observations clearly show a majority of non-mineralized cells alongside the remaining, cell-free, mineralized S-layer shells. Analyses of ^15^N uptake using NanoSIMS confirmed this timing, showing no significant ^15^N incorporation for mineralized samples prior to 2 d of recovery in labelled medium, compared to T0 control aliquots that were removed immediately after resuspension of cells in the labelled medium (see Fig. 6). The S-layers of mineralized samples remained at a relatively steady ^15^N/(^14^N + ^15^N) ratio (see green regions in Supplementary Figure 6, mineralized samples) compared to the emerging cells, showing that mineralized S-layers did not incorporate ^15^N during the cell division processes giving rise to the emerging cells. This is coherent with S-layer shedding and complete regeneration. S-layer presence before and after mineralization and shedding was confirmed by SDS-PAGE and mass spectrometry (see Supplementary Figure 4 and 5).

## 4. Discussion

### 4.1 *Lysinibacillus* sp. TchIII 20n38 Is Highly Adapted for Survival Under Stress Conditions

The bacterial isolate used in this study, *Lysinibacillus* sp. TchIII 20n38, is a metallotolerant grampositive bacterium (*article in preparation*)., possessing a cell envelope composed of the plasma membrane surrounded by a thick layer of peptidoglycan capped by an S-layer forming an ordered structure at the cell surface. This flexible cage-like structure is in direct contact with the surrounding environment, and thus provides the primary protective element against potentially toxic environmental concentrations of heavy metals or radionuclides. *Lysinibacillus* sp. TchIII 20n38 demonstrated many adaptive mechanisms to the stresses induced by nutrient limitation and/or the presence of iron, as might be expected for a bacterium isolated from a radioactive waste disposal site. These included the accumulation and enlargement of intracellular polyphosphate granules and sporulation under phosphate-limiting conditions, cell size reduction and morphology alterations from rod-shaped to ovoid cells after metal stress, as well as reductions in biofilm after exposure to either iron or acidic pH and augmentation in biofilm after a return to neutral pH in the absence of additional iron input. Polyphosphate accumulation is a common mechanism used by bacteria, including *Lysinibacillus sphaericus*, in response to nutrient stress (depletion of amino acids), and prior to sporulation (Shi et al., 2015; Tocheva et al., 2013).

### 4.2 S-layer Shedding Mechanism in *Lysinibacillus* as a Response to Metal Stress

S-layers from a variety of prokaryotes are known to induce mineral formation. The S-layers of cyanobacteria are able to nucleate selenite and strontium (Schultze-Lam and Beveridge, 1994), while the S-layers of thermophilic archaea can form amorphous Fe-phosphate minerals in the quasi- periplasmic space between the S-layer and the underlying lipid membrane (Kish et al., 2016). Diverse *Lysinibacillus* sp. have been observed to precipitate minerals on their cell surfaces, including U- phosphates (Merroun et al., 2005; Mondani et al., 2011) and calcium carbonate (Kaur and Mukherjee, 2013). Here, we show that *Lysinibacillus* sp. TchIII 20n38 cells become encrusted with Fe-minerals after exposure to high concentrations of iron under mildly acidic conditions. Mineral precipitation by the cell surfaces, including S-layers, prevent damages to cells including oxidative stress, enzyme deactivation, protein denaturation, and membrane disruption (Lemire et al., 2013). Iron precipitation nucleated by the S-layer proteins prevents an overproduction of free radicals in the cytosol due to Fenton chemistry.

While mineral formation on S-layers is known, the mechanisms for removing such barriers to exchange with the surrounding environment are not as well understood. Mechanisms identified to date have described partial removal of cell envelope components after metal interactions, particularly membrane vesicle formation (Kish et al., 2016; McBroom and Kuehn, 2007; Shao et al., 2014). Partial shedding of S-layer fragments has also been observed, both for mineralized cyanobacterial Slayers (Schultze-Lam and Beveridge, 1994; Schultze-Lam et al., 1992) and non-mineralized S-layers for stationary phase bacteria likely as part of cell wall turnover (Luckevich and Beveridge, 1989). During the course of normal cell growth in the closely related *Lysinibacillus sphaericus*, bands of Slayer monomer insertion form on cell surfaces, and in the course of cell division new S-layer monomers are only inserted at the newly-formed the poles (Howard et al., 1982).

Here we show that *Lysinibacillus* sp. cells were able to recover normal growth after mineral encrustation through a shedding of mineralized S-layers, followed by S-layer regeneration. To our knowledge, this is the first report of complete S-layer shedding and regeneration. S-layer shedding required an additional 24h before cells returned to normal cell division compared to non-mineralized cells, as shown by the clustering of ^15^N uptake by 2 d mineralized cells with 1 d non-mineralized samples as measured by NanoSIMS over a time course after exposure to iron. Uptake of ^15^N also illustrated that despite extensive mineralization, cells retained active metabolism. The continued presence of shed, mineralized S-layers composed of ^14^N-based proteins alongside cells bearing newly regenerated ^15^N-bearing S-layers resulted in the heterogeneity in ^15^N-incorporation for the M_2d samples (see Fig. 6). S-layer shedding activity was limited to cells in mid-exponential growth phase, providing an advantage over cells in stationary growth phase in Fe-rich conditions.

### 4.3 Potential Applications of S-layer Regeneration in Bioremediation

S-layers have a strong potential for use biotechnology, due to the combination their capacities for both self-assembly and metal binding. S-layers have been used as templates to nucleate the fabrication of metal nanoparticles (Pollmann and Matys, 2007) that can be applied as biocatalysts(Creamer et al., 2007) or exploited for their unique magnetic properties (Bartolomé et al., 2012). S-layers assembled in vitro have been proposed as bio-filters for bioremediation technologies(Pollmann et al., 2006).

The total S-layer shedding and regeneration shown here after mineral encrustation opens further possibilities for a self-renewing bioremediation system, employing soil bacteria already tolerant to metals and radionuclides.

## 6 Author Contributions

AK and LR contributed conception and design of the study; AC performed the experiments, LR performed NanoSIMS measurements, CL and SZ performed mass spectrometry analyses; ED performed the statistical analysis of NanoSIMS data; AK wrote the first draft of the manuscript; AC wrote sections of the manuscript. All authors contributed to manuscript revision, read and approved the submitted version.

## 8 Acknowledgments

The authors gratefully acknowledge Virginie Chapon for giving us the strain used in this study, along with helpful discussions. The authors thank Adriana Gonzalez-Cano and Rémi Duhamel for assistance with NanoSIMS analyses, and Imène Esteve of the Scanning Electron Microscope (SEM) facility of the Institut de Minéralogie, Physique des Matériaux et Cosmochimie for assistance with imaging, which was greatly appreciated. We also wish to thank Jennyfer Miot for productive discussions.

## 10 Supplementary Material

The Supplementary Material for this article can be found as a separate file.

